# Exploring the Pharmacological Mechanism of Bezoar on Cerebral Ischemic Stroke using a Network Pharmacology Approach

**DOI:** 10.1101/2020.02.28.969436

**Authors:** Xin Du, Changxiang Li, Beida Ren, Nan Deng, Congai Chen, Xueqian Wang, Fafeng Cheng, Min Liu Qingguo Wang

## Abstract

**Background:** Ischemic stroke is a common clinical disease limited by its time window for treatment. In addition to its high mortality rate, only one treatment currently exists for ischemic stroke. Nonetheless, traditional Chinese medicine is often used as a reatment for the disease. Bezoar is a multi-functional drug that has been demonstrated to be effective for the treatment of ischemic stroke. However, its mechanism of action is yet to be fully elucidated. Based on network pharmacology, we explored the potential mechanism of action of bezoar. Symmap and literature data mining methods were used to obtain the target of bezoar. The mechanism of bezoar for the treatment of ischemic stroke was identified and ischemic stroke-related targets were retrieved using DrugBank, Online Mendelian Inheritance in Man, and Therapeutic Target Database. Protein-protein interaction networks were constructed using the Cytoscape plugin, BisoGenet, and analysed by topological methods. Gene Ontology and Kyoto Encyclopaedia of Genes and Genomes pathway enrichment were carried out via the Database for Annotation, Visualization, and Integrated Discovery server. We obtained 48 potential targets and 3 signalling pathways, including mitogen-activated protein kinase, hypoxia-inducible factor-1, and tumour necrosis factor signalling pathways. The mechanism of action of bezoar on ischemic stroke involves multiple targets and signalling pathways. Our research provides a network pharmacology framework for future Chinese medicinal research.

## 1. Introduction

Cerebral ischaemia is a major cause of disability and death globally. In addition, its most common manifestation is stroke. In fact, 85% of stroke cases are ischaemic and it is the second leading cause of death and most common cause of complex chronic disability worldwide.^[1]^ Ischemic stroke results in the reduction of blood flow from the occluded artery to the brain tissue.^[2]^ Currently, thrombolytic therapy is the main treatment for ischemic stroke. However, thrombolytic therapy is limited by its narrow effective time window and the risk of bleeding.^[3]^Therefore, potential drugs for ischemic stroke treatment are urgently needed. The pathological process of cerebral ischemia–reperfusion injury is complex and unclear. In addition, it involves multiple mechanisms, such as damage and inflammation caused by free radicals, energy metabolism disorder in brain tissues, toxicity of excitatory amino acids, overload of intracellular calcium, cytotoxic effect of NO, and abnormal opening of the blood–brain barrier.^[4]^To resolve these problems, future drug research should focus on combination therapy and multi-target intervention.

Traditional Chinese medicine has been demonstrated to be effective in the treatment of ischemic stroke, with different effects in the acute and convalescent periods. Refined Qingkailing, a clinical treatment widely used to treat ischemic stroke, reduces the infarct size of cerebral ischemia and increases the expression levels of endothelial nitric oxide synthase after middle cerebral artery occlusion (MCAO).^[5]^Bezoar contains taurine, bile acid, bilirubin, and tauroursodeoxycholic acid (TUDCA). Taurine has antioxidative properties, stabilises the membrane, functions as an osmoregulator, modulates ionic movements, reduces the level of proinflammatory cytokines, and regulates the intracellular calcium concentration, which contribute to its neuroprotective effect.^[6]^TUDCA activates the Akt-1/protein kinase B survival pathway and induces BCL2 associated agonist of cell death phosphorylation at Ser-136 to exert neuroprotective effects.^[7]^TUDCA also influences the microglial phenotype in vivo and in vitro toward the anti-inflammatory. G protein-coupled bile acid receptor 1 may be a new therapeutic target for pathologies associated with neuroinflammation and microglial activation, such as traumatic brain injuries, stroke, and neurodegenerative diseases.^[8]^

Different components of bezoar exhibit a therapeutic effect on ischemic stroke; however, the therapeutic mechanism remains at the animal level and is insufficient to demonstrate its good efficacy. Therefore, the development of modern and technological approaches is urgently needed to analyse the action mechanisms of bezoar in the treatment of various diseases. Fortunately, modern technologies and medical sciences have rapidly promoted the development of the systems pharmacology method, which is a promising strategy to elucidate the molecular mechanisms of drugs. By using a systematic approach in network pharmacology, the relationship between drugs, targets, and diseases can be visually presented in a drug target network.^[9]^In this study, we used a network pharmacology platform^[10]^ for drug target prediction, topology screening, and gene functional analysis to uncover the mechanism of bezoar on ischemic stroke and show its medicinal value. Additionally, we used a novel research method to elucidate the mechanistic effects of bezoar in the treatment of ischemic stroke and identify potential protein targets.

In this study, we employed network pharmacology to explore the mechanism used by bezoar when used for the treatment of stroke. Briefly, as shown in Figure 1, we first collected information on bezoar from TCMSP, Symmap, and Pubchem and screened for eligible compounds using ADME screening. Subsequently, we sought to predict the relevant protein targets and target information of related diseases using OMIM, TTD, Drug bank, Gene Cards, and CTD and construct the compound information network using KEGG and GO analyses.

**Fig 1:**
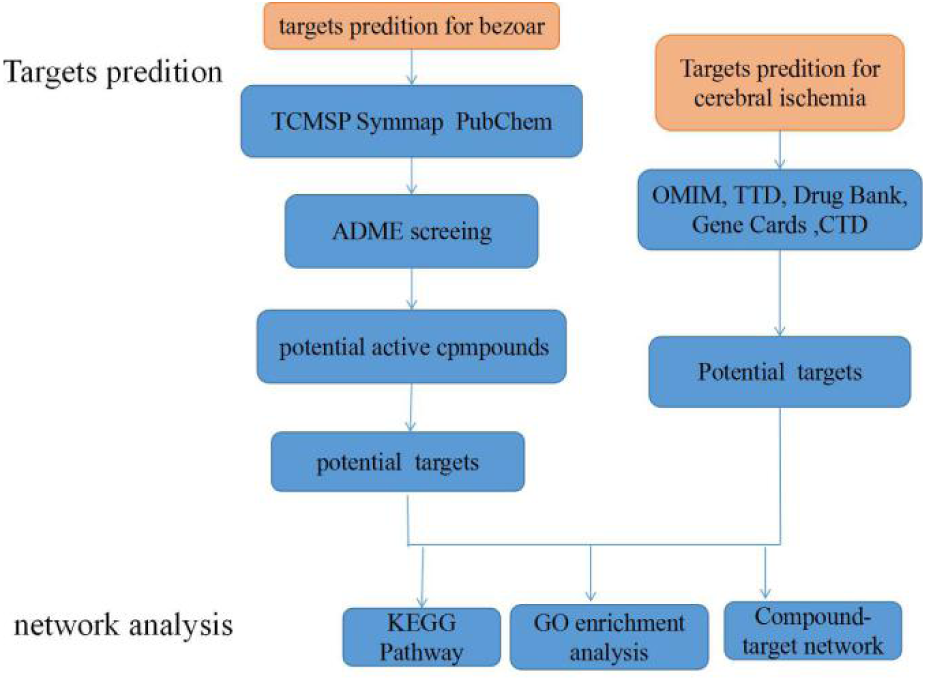
System pharmacology flow chart.

## 2 Methods

### 2.1 Aim of the study

Network pharmacology was used to explore the mechanism of bezoar in the treatment of stroke.

### 2.2 Components of bezoar

All compounds in bezoar were collected using a wide-scale text mining approach. Databases such as the Traditional Chinese Medicine Systems Pharmacology (TCMSP) database (http://lsp.nwu.edu.cn/tcmsp.php) and Symmap (http://www.symmap.org/) were used. Structural information of the components was obtained from the National Center for Biotechnology Information (NCBI) PubChem (https://pubchem.ncbi.nlm.nih.gov/).

### 2.3 Potential active compound screening

Absorption, distribution, metabolism, and excretion (ADME) evaluations are critical procedures for the screening of potential active drugs with reasonable pharmacokinetic and pharmacological properties. Oral bioavailability (OB) represents the rate and extent to which the active ingredient or moiety is absorbed from the drug and available at the site of action.^[11]^In this study, an ADME system that integrated the predicted OB (PreOB) and predicted drug-likeness (PreDL) model was employed for bioactive ingredient screening. Compounds with OB > 30% and DL > 0.18 were selected as potential active compounds and used for further studies.

### 2.4 Target recognition of bezoar components

To determine the potential targets of bezoar, target prediction of the bezoar components was performed. Chemical structures were obtained from PubMed Central in the NCBI database. To retrieve the targets for the compounds, different databases, such as Stitch(http://stitch.embl.de/), Swiss Target Prediction (http://swisstargetprediction.ch/), Sea (http://sea.bkslab.org), and Target Net (http://targetnet.scbdd.com/) were searched.

### 2.5 Predicting ischemic stroke-associated targets

The Therapeutic Target Database (TTD https://db.idrblab.org/ttd/)^[12]^,Online Mendelian Inheritance in Man (OMIM https://omim.org/)^[13]^, Drug Bank (https://www.drugbank.ca/)^[14]^, Gene Cards (https://www.genecards.org/), and Comparative Toxicogenomics Database (CTD http://ctdbase.org/)^[15]^ were searched with the keywords “cerebral ischemic stroke”, “cerebral ischemia”, and “cerebral infarction”.

### 2.6 Screening for key target proteins

A Venn diagram was used to identify the targets of the drug compounds and the disease to show the key targets for drugs to treat ischemic stroke. The Venn diagram was plotted using the OmicShare platform, a free online platform for data analysis (http://www.omicshare.com/tools/)^[16]^.

### 2.7 Protein-protein interaction (PPI) data

Data on PPIs were obtained from STRING (https://string-db.org/; version 10)^[17]^, with the species limited to “homo sapiens” and a confidence score > 0.4. PPI networks were constructed using the PPI network derived from STRING. The results were analysed using the Cytoscape database.

### 2.8 Gene Ontology (GO) and Kyoto Encyclopaedia of Genes and Genomes (KEGG) pathway enrichment analysis

GO analysis is widely used for the annotation of gene function. It consists of three parts: biological process, cellular component, and molecular function. In contrast, KEGG is a database of integrated chemistry, genomic, and systems function information that is used to understand the advanced functions and uses of biological systems from molecular level information. Based on the gene annotation function of the GO database and the access information of the KEGG database, The Database for Annotation, Visualization and Integrated Discovery (DAVID https://david.ncifcrf.gov/) provides a systematic bioinformatics annotation method for large-scale genes and proteins and provides enrichment analysis. The key targets of bezoar in ischemic stroke was uploaded into the DAVID database. Gene function and pathway enrichment analyses were performed on the key targets of bezoar on haemorrhagic stroke. A P value ≤ 0.05 was considered significant. GO enrichment analysis was also used to perform a hypergeometric test.

## 3 Results

### 3.1 Bezoar components

Using Symmap and TCMSP, we derived important information related to ingredients, targets, and the disease. We obtained 48 ingredients and 134 related targets from Symmap and 19 ingredients and 133 related targets from TCMSP; hence, we obtained a total of 277 compound targets.

### 3.2 Potential active compound and ischemic stroke targets

Using TCMSP and Symmap, information related to ADME, such as human OB, DL, Caco-2 permeability, blood-brain-barrier (BBB) permeability, and Lipinski’s rule of five (based on molecular weight, AlogP, topological polar surface area, H bond donors, and H bond acceptors) was derived. Based on the principle that OB > 30% and DL > 0.18, we retrieved 8 compounds (Table 1).

**Table 1:** Pharmacokinetic parameters of candidate active compounds in bezoar

| Drug ID | Molecular name | CAS number | OB (%) | DL (%) |
| --- | --- | --- | --- | --- |
| SMIT00540 | Belamcandal | 138501-57-2 | 30.0718 | 0.67 |
| SMIT01395 | Embinin | 52589-13-6 | 33.91 | 0.73 |
| SMIT03442 | Clr | 80356-14-5 | 37.8739 | 0.68 |
| SMIT10058 | Methyltetradecahydro<br>pentanoate | 1448-36-8 | 32.3177 | 0.76 |
| SMIT10059 | Methyldeoxycholate | 3245-38-3 | 34.6346 | 0.73 |
| SMIT10064 | Deoxycholic acid | 83-44-3 | 40.723 | 0.68 |
| SMIT10065 | Zinc01280365 | 64-85-7 | 46.3756 | 0.49 |
| SMIT19325 | Mangiferolic acid | 4184-34-3 | 36.16 | 0.84 |

Using five available databases, TTD, OMIM, DrugBank, Gene Cards, and CTD, we obtained 32, 152, 90, 63, and 118 ischemic stroke-related targets, respectively. We removed duplicate targets and obtained a total of 384 targets (Figure 2).

**Fig 2:**
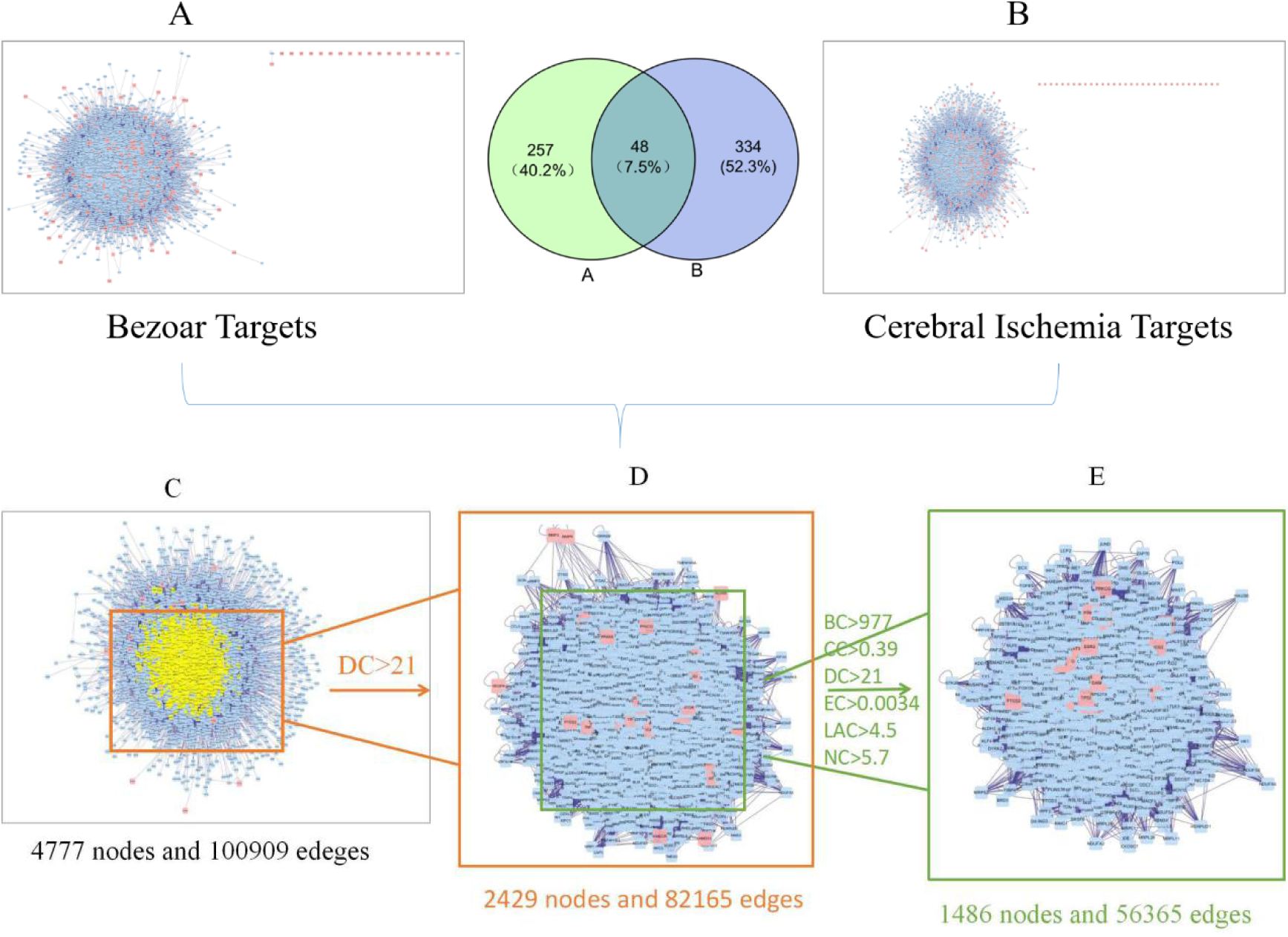
Core PPI network of bezoar against ischemic stroke. (A) PPI network for the bezoar target. (B) PPI network for ischemic stroke targets. (C) An interactive PPI network of bezoar and ischemic stroke targets, including 4,777 nodes and 100,909 edge. (D) PPI network of important proteins extracted from (c); the network consists of 2,429 nodes and 82,165 edges. (E) The important protein extracted from (d) by the PPI network; the network consists of 1,486 nodes and 56,365 edges. BC: Intermediary Centre; CC: Close to Centre Status; DC: Degree Centre; EC: Feature Vector Centre; NC: Network Centre; LAC: Local Average Connection.

Thereafter, we constructed the bezoar target network and an ischemic stroke-related target network using PPI data (Figure 2A and B). To reveal the pharmacological mechanism of bezoar for ischemic stroke, we used a combined network function. This new network (4,777 nodes and 100,909 edges), which was built from two networks (Figure 3C) with overlapping targets, was provided by Cytoscape. As nodes that have degrees amounting to only twice the median of all nodes can serve as important targets, we constructed a meaningful network of 2,429 nodes and 82,165 edges (Figure 3D) for bezoar targets in ischemic stroke. Finally, we selected six topological features to confirm the candidate targets, Intermediary Centre (BC), Close to Centre Status (CC), Degree Centre (DC), Feature Vector Centre (EC), Network Centre (NC), and Local Average Connection (LAC), with CytoNCA. The candidate targets showed values of BC > 977, CC > 0.39, DC > 21, EC > 0.0034, NC > 5.7, and LAC > 4.5 (Figure 3E).

**Fig 3:**
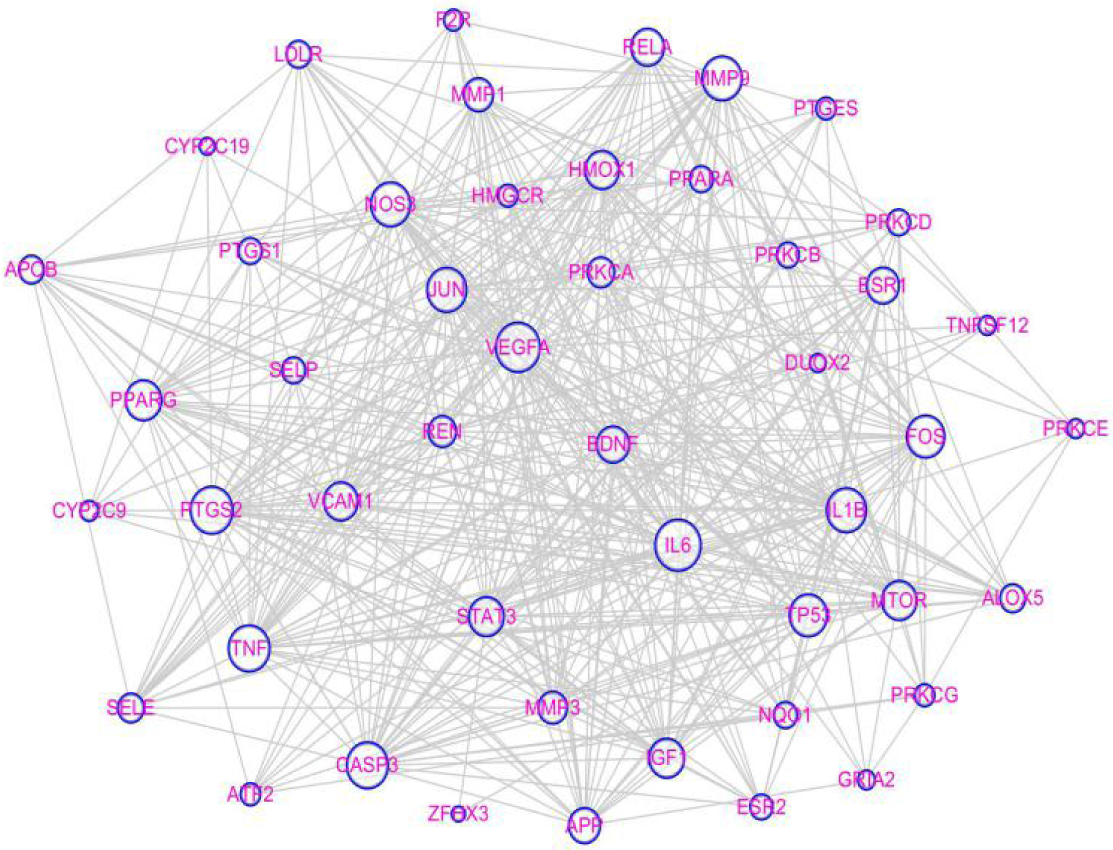
Direct interaction between the 48 compound targets and ischemic stroke proteins. Node size is proportional to its degree.

### 3.3 Key targets for drugs and diseases

By using a Venn diagram, we identified 48 common targets of drugs and diseases (Table 2).

**Table 2:**
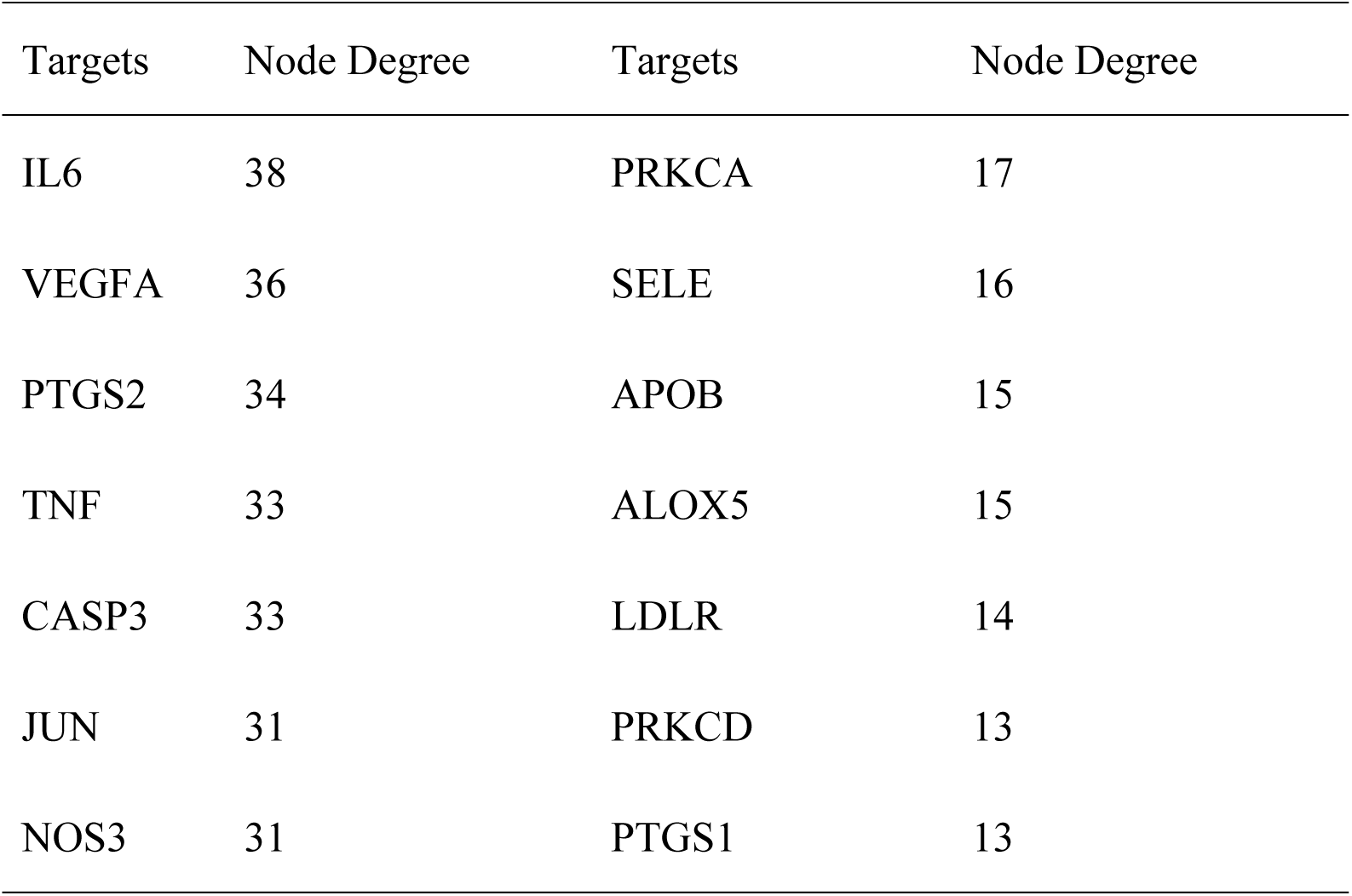
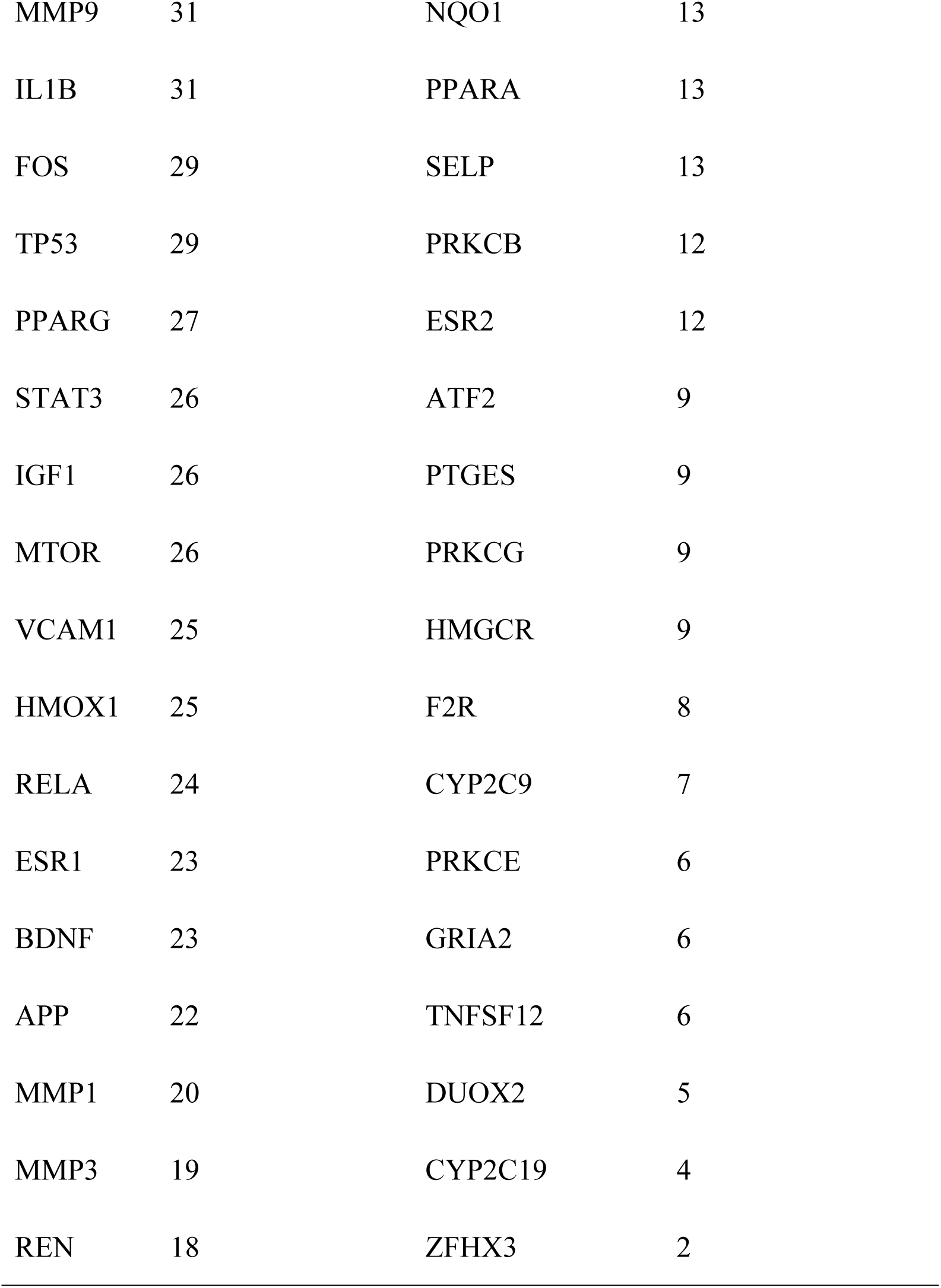
Node degree of the targets of Bezoar targets via Cytoscape

| Targets | Node Degree | Targets | Node Degree |
| --- | --- | --- | --- |
| IL6 | 38 | PRKCA | 17 |
| VEGFA | 36 | SELE | 16 |
| PTGS2 | 34 | APOB | 15 |
| TNF | 33 | ALOX5 | 15 |
| CASP3 | 33 | LDLR | 14 |
| JUN | 31 | PRKCD | 13 |
| NOS3 | 31 | PTGS1 | 13 |
| MMP9 | 31 | NQO1 | 13 |
| IL1B | 31 | PPARA | 13 |
| FOS | 29 | SELP | 13 |
| TP53 | 29 | PRKCB | 12 |
| PPARG | 27 | ESR2 | 12 |
| STAT3 | 26 | ATF2 | 9 |
| IGF1 | 26 | PTGES | 9 |
| MTOR | 26 | PRKCG | 9 |
| VCAM1 | 25 | HMGCR | 9 |
| HMOX1 | 25 | F2R | 8 |
| RELA | 24 | CYP2C9 | 7 |
| ESR1 | 23 | PRKCE | 6 |
| BDNF | 23 | GRIA2 | 6 |
| APP | 22 | TNFSF12 | 6 |
| MMP1 | 20 | DUOX2 | 5 |
| MMP3 | 19 | CYP2C19 | 4 |
| REN | 18 | ZFHX3 | 2 |

### 3.4 Construction of the target protein interaction network

We used String and Cytoscape to construct a protein interaction network for the molecular mechanism using 48 nodes and 453 edges. Removal of these inclusions would result in crashing of the network (Figure 3).

### 3.5 GO enrichment analysis of targets for bezoar against ischemic stroke

First, we performed GO analysis of bezoar for the 48 candidate targets for ischemic stroke. Thereafter, DAVID was used to analyse the biological functional units and potential biological system functions. The results were divided into biological process, cellular component, and molecular function (Figure 4). The biological process was related to positive regulation of transcription, DNA-templating, positive regulation of smooth muscle cell proliferation, positive regulation of nitric oxide biosynthetic process, positive regulation of transcription from RNA polymerase II promoter, aging, positive regulation of sequence-specific DNA binding transcription factor activity, platelet activation, leukocyte tethering or rolling, response to cAMP, and response to drug.

**Fig 4:**
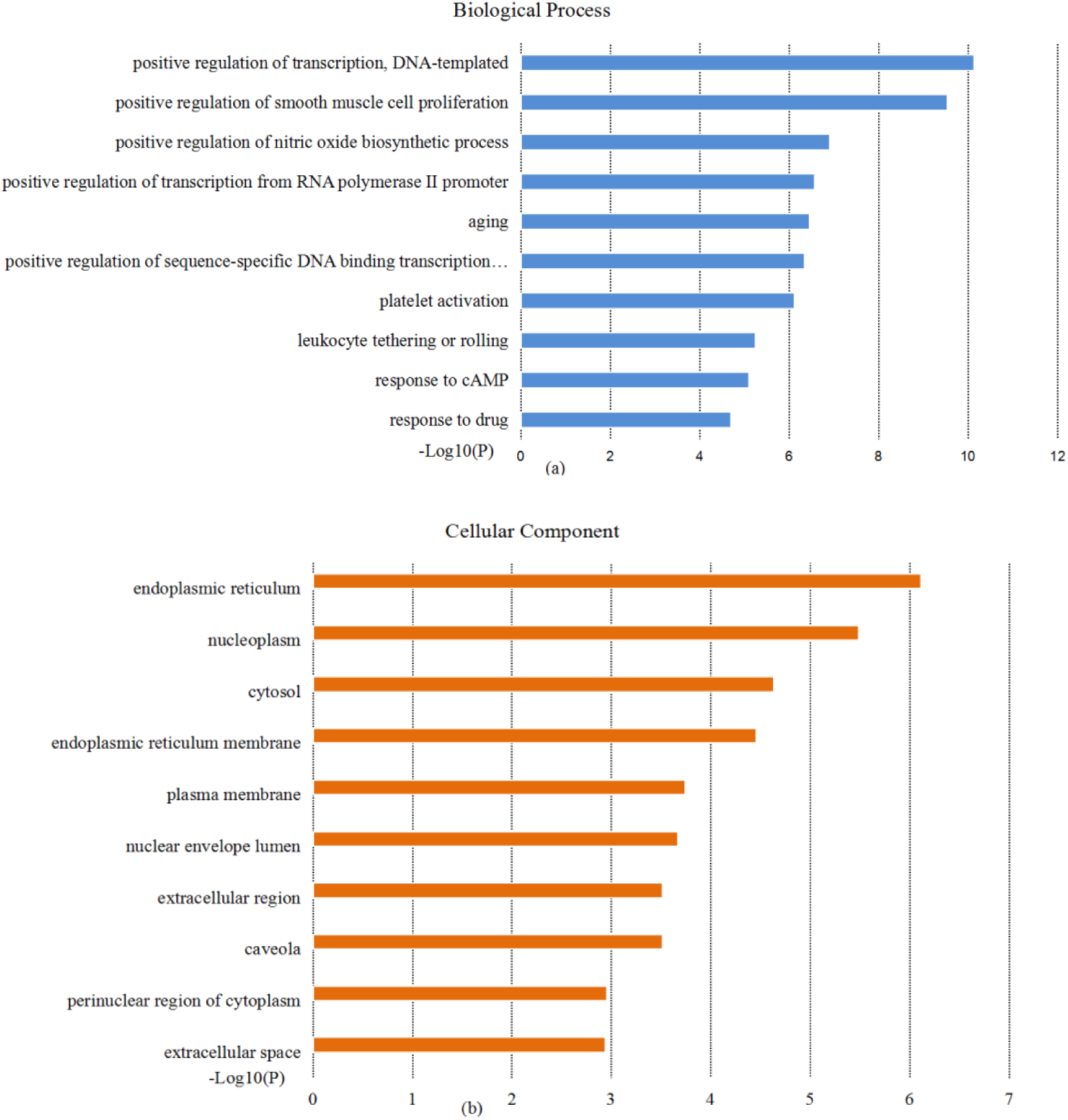
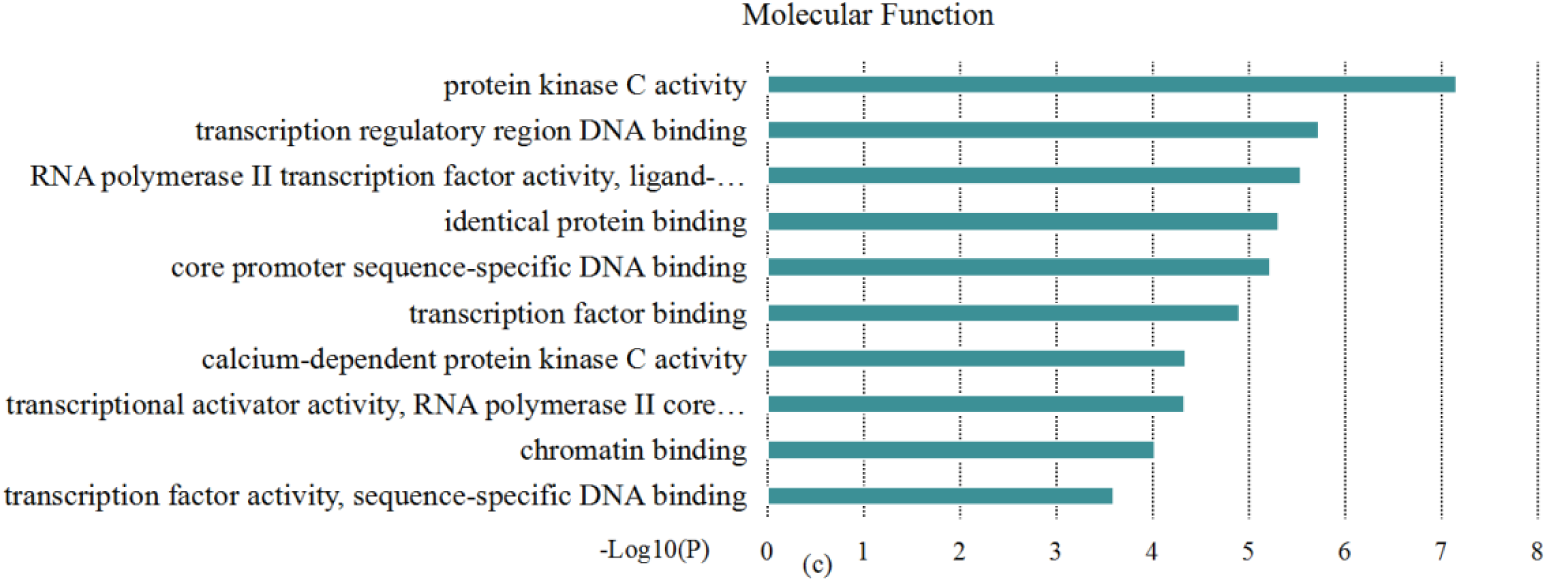
GO analysis was performed on the screened genes. The top 10 terms for (a) biological processes, (b) cell component, and (c) molecular function with P < 0.05 are shown.

The cellular component was related to extracellular space, perinuclear region of the cytoplasm, caveola, extracellular region, nuclear envelope lumen, plasma membrane, endoplasmic reticulum membrane, cytosol, nucleoplasm, and endoplasmic reticulum. The molecular function was related to protein kinase C activity, transcription regulatory region, DNA binding, RNA polymerase II transcription factor activity, ligand-activated sequence-specific DNA binding, identical protein binding, core promoter sequence-specific DNA binding, transcription factor binding, calcium-dependent protein kinase C activity, transcriptional activator activity, RNA polymerase II core promoter proximal region sequence-specific binding, chromatin binding, transcription factor activity, and sequence-specific DNA binding.

### 3.6 GO and KEGG pathway enrichment analysis

To elucidate the important role of bezoar targets in the treatment of ischemic encephalopathy, we used a comprehensive ischemic stroke approach based on current knowledge of the pathogenesis of ischemic stroke. The top 10 KEGG signalling pathways for bezoar were obtained and constructed with P values (Figure 5). Based on this system-level image, mitogen-activated protein kinase (MAPK), hypoxia-inducible factor-1 (HIF-1), and tumour necrosis factor (TNF)-α were selected.

**Fig 5:**
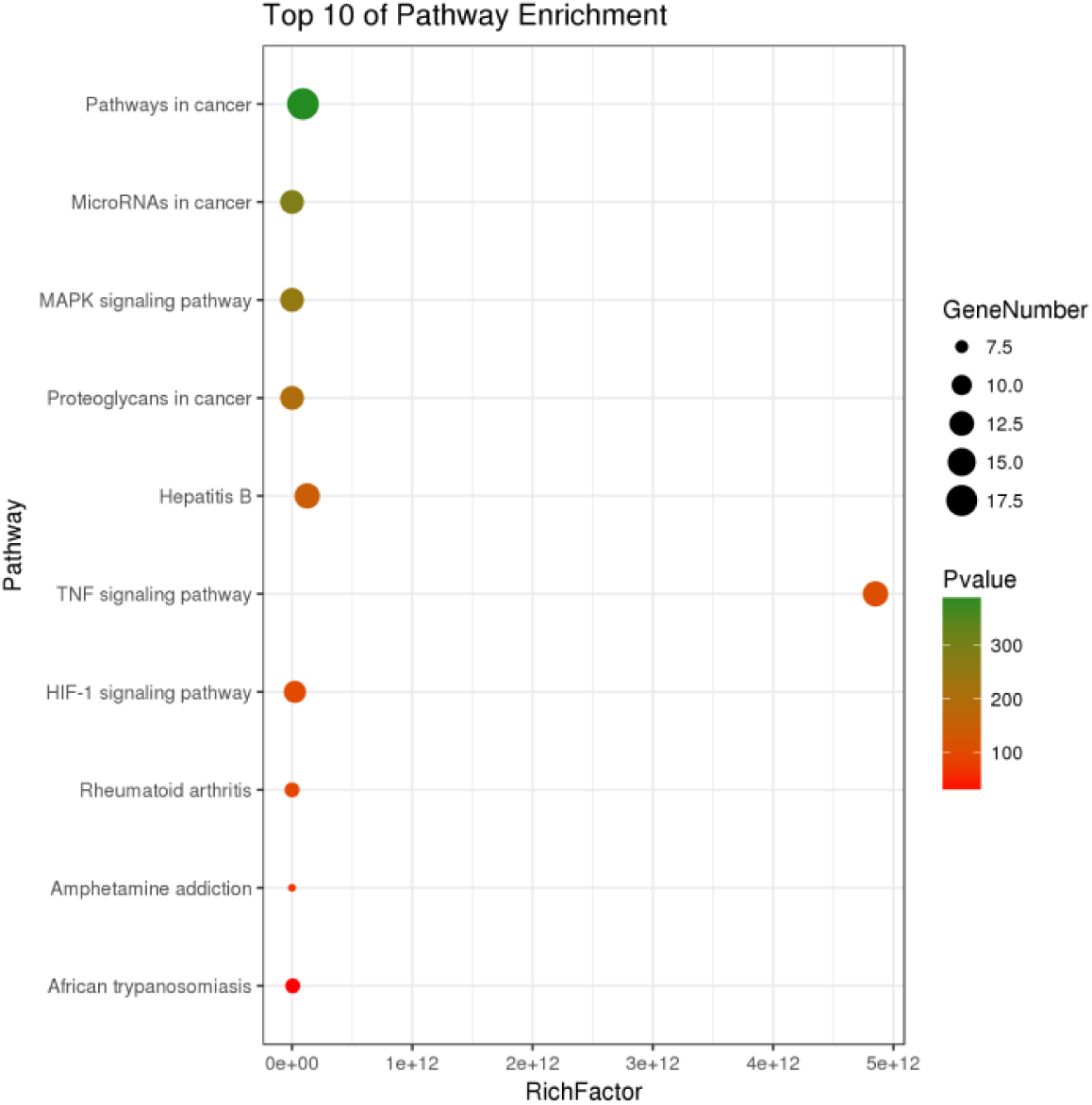
KEGG signalling pathway is enriched for the screening of genes. “Rich factor” represents the proportion of the number of target genes, number of pathways, and annotated genes located in the pathway. Higher rich factors indicate higher levels of enrichment. The size of the dots indicates the number of target genes in the pathway and the colour of the dots reflects different P values.

## 4. Discussion

Network pharmacology assists in the identification of drug targets, mimics signal pathways for drug and disease interactions, improves clinical outcomes, and reduces laboratory costs.^[18]^Herb and plant ingredients have positive prospects for the treatment of complex diseases such as ischemic stroke because of the few side effects and multiple targets.^[19]^We used network pharmacology-related methods to elucidate the therapeutic targets and mechanistic pathways of bezoar for ischemic stroke.

TUDCA can alleviate acute brain damage caused by cerebral ischemia, prevent apoptosis, protect mitochondrial integrity, and inhibit caspase activation.^[20]^We showed that taurine can exert a protective function against hypoxia by increasing cell viability, decreasing infarct volume, and reducing endoplasmic reticulum stress in vitro and in vivo. Taurine can decrease apoptosis by the downregulation of apoptotic markers and caspase-12 in primary neuronal culture.^[21]^

In this study, we identified 383 disease targets and 277 drug targets. By constructing a Venn diagram, we obtained 48 common targets for diseases and drugs and subsequently used the PPI network to retrieve a map of the interactions of the 48 targets. Using a topological approach, we identified a core network containing 48 nodes and 453 edges. GO analysis suggested that bezoar plays an important role in the treatment of ischemic stroke, mainly via the positive regulation of transcription, DNA-templating, extracellular space, and transcription regulatory region DNA binding. This is also suggested by KEGG analysis of the core PPI network. Hence, bezoar may have multiple effects on stroke treatment by regulating multiple pathways. Based on each P-rich pathway and their relationship with ischemic stroke, three signalling pathways, MAPK, TNF, and HIF-1, were most concerning. Hence, we constructed a signal pathway map to elucidate the potential molecular mechanism of bezoar in ischemic stroke.

In this analysis, based on pathway enrichment analysis and our current understanding of ischemic stroke, the pathways directly related to the pathological process of cerebral ischemia were included in the “integration pathway for cerebral ischemia”. Integration pathways include the MAPK, HIF-1, and TNF-α signalling pathways.

The activation of the nuclear factor (NF)-κB or MAPK signalling pathway is partly responsible for the induced expression and activation of nucleotide-binding oligomerization domain, leucine rich repeat and pyrin domain containing protein (NLRP)1 and NLRP3 inflammasome proteins and such effects can be attenuated using pharmacological inhibitors of these two pathways in neurons and brain tissue under in vitro and in vivo ischemic conditions, respectively.^[22]^Taurine exerts potent antithrombotic effects by reducing intravascular fibrin and platelet deposition, which in turn reduces downstream microvascular thrombosis and improves cerebral vascular potency. Inhibition of NF-κB and CD147 may be involved in the mechanisms of microvascular protection by taurine. Taurine inhibits NF-κB activation in the brain after ischemic stroke.^[23]^P38 MAPK is related to the activation and phosphorylation of cPLA2 and arachidonic acid release.^[24]^The interaction between 12/15-LOX and p38 MAPK/cPLA2 pathways promotes the progression of arachidonic acid metabolism and exacerbates the inflammatory process and tissue injury.^[25]^MAPK consists of three major members, p38, extracellular signal-regulated kinase 1/2 (ERK1/2), and c-Jun N-terminal kinase (JNK), which direct extracellular signals to intracellular targets to regulate cellular activity through various signalling pathways.^[26]^MAPKs, as regulators of NF-κB, play an important role in inflammatory factors.^[27]^

NF-κB is an early nuclear transcription factor, whereas NF-κB p65 is an important member of the NF-κB transcription family and plays a central role in inflammatory responses. Under normal physiological conditions, NF-κB is present in the cytoplasm in an inactive form and is activated when stimulated by external antigens. It is also transferred to the nucleus to regulate the expression of downstream inflammatory factors. Inflammation regulated by HIF-1α can be mediated by nuclear translocation of pyruvate kinase (PK)M2. However, NF-κB regulates the expression of HIF-1α and activation of NF-κB p65 is positively correlated with the expression of HIF-1α.^[28]^The NF-κB signalling pathway regulates inflammation by promoting the expression of interleukin (IL)-1β, IL-6, TNF-α, and other proinflammatory cytokines.^[29]^P38 MAPK is involved in the mitochondrial kinetics of cerebral ischemic injury, which indicates the potential neuroprotective effects of p38 inhibition.^[30]^

HIF-1 is a key regulator in hypoxia that is composed of HIF-1α and HIF-1β protein subunits.^[31]^HIF activity is primarily affected by oxygen-dependent regulation of the HIF-κ subunit protein amount through the HIF prolyl 4-hydroxylase domain (PHD) protein family. PHD activity is dependent on the availability of molecular oxygen, ferrous iron, and 2-oxoglutarate.^[32]^The activity of HIF-1 is mainly determined by the level of the alpha subunit and is involved in cerebrovascular diseases under various pathological conditions, such as ischemic stroke and traumatic brain injury. In adult and neonatal rodent ischemic stroke models, inhibition of HIF-1 ameliorates hypoxia-induced BBB destruction and subsequent brain damage. HIF-1 disrupts the integrity of the BBB by enhancing the expression of its target gene, vascular endothelial growth factor (VEGF).^[33]^HIF-1α is a classic activator of VEGF production. The HIF-1α/VEGF pathway plays an important role in neurorepair and functional recovery following experimental stroke.^[34]^HIF-1 is generated in the brain in response to ischemia or hypoxia and its activity is determined by the availability and activity of the oxygen-regulated subunit, HIF-1α.^[35]^HIF-1 exerts its activity through proteins encoded by its downstream genes, such as erythropoietin, VEGF, and glucose transporters.^[36]^VEGF is a pleiotropic growth factor that plays an important role in neurogenesis, axonal plasticity, angiogenesis, and vascular permeability. Its expression increases in the ischemic penumbra within 3 h after ischemia. As a downstream product of HIF-1, VEGF can induce vascular leakage and matrix metallopeptidase 9 (MMP-9) activation.^[37]^The early administration of VEGF to ischemic rats can enhance BBB leakage and increase the area of infarct in the brain.^[38]^Inflammatory responses play an important role in brain ischemia/reperfusion (I/R) injury. In addition, inflammation might be one of the mechanisms of BBB destruction. HIF-1α plays an important role in the inflammatory process. VEGF, an important downstream gene of HIF-1α, is closely related to BBB leakage in acute ischemic brain. The accumulation of HIF-1α and subsequent upregulation of VEGF contribute to basement membrane damage and enhance BBB leakage, ultimately exacerbating hyperosmotic brain oedema.^[39]^Signal transduction and transcription activator-3 (STAT3) is a member of the STAT protein family of transcription factors that coordinate and integrate signals from extracellular stimuli and play an important role in the growth and differentiation of a variety of cells.^[40]^There is evidence that cerebral ischemia promotes STAT3 activation and nuclear translocation as well as STAT3-dependent transcription of target genes, including HIF-1α.^[41]^ STAT3 is a potential regulator of HIF-1α-mediated VEGF.^[42]^ STAT3 also plays an important role in neuronal survival and regeneration. Cerebral ischemia promotes STAT3 activation and nuclear translocation as well as STAT3-dependent transcription of target genes, including HIF-1α.^[43]^During ischemic stroke, a decrease in oxygen levels activates HIF and this induces the expression of cytoprotective factors, such as VEGF and erythropoietin.^[44]^

The MAPK family includes ERK1/2, JNK, and p38. MAPK is activated by phosphorylation, causes target gene expression, and its signalling pathway is involved in the regulation of inflammatory response, cytokine expression, and apoptosis after stroke.^[45]^ Activation of the MAPK signalling pathway in ischemic stroke is important and ERK and JNK activation leads to a significant reduction in inflammatory cytokines and apoptosis in the injured area.^[46]^MAPK can be activated by phosphorylation of threonine and tyrosine residues in response to cerebral ischemia. When phosphorylated, it supports neuronal survival in the dentate gyrus. Hence, the EGFR/MAPK signalling cascade is an important therapeutic target for inducing neurogenesis and brain repair for post-stroke disability.^[47]^P38 MAPK plays a dual role in the regulation of cell death. Its activation after focal cerebral ischemia promotes the production of pro-inflammatory cytokines and aggravates infarction.^[48]^The phosphorylation of p38 MAPK activates cerebral deficits by activating anti-apoptotic signals in the ischemic cortex. Blood damage plays a neuroprotective role in hypoxic preconditioning and the transient MCAO model. Pharmacological activation of the PI3K/Akt signalling pathway plays an anti-apoptotic role in ischemic penumbra after brain I/R injury.^[49]^

NF-κB and MAPK signalling cascades, including ERK1/2, JNK, and p38 activation, in the cerebral ischemic response plays a key role in regulating proinflammatory mediator production and apoptosis.^[50]^ The MAPK pathway contributes to hypoxia-induced apoptosis in neuroblastoma cells. Hypoxia activates the MAPK signalling pathway by enhancing phospho-ERK, JNK, and p38 MAPK in different cells.^[51]^ Different signalling pathways activate different transcriptional MAPK factors, mediating different biological effects, but there are also extensive cross-links in these pathways and phosphorylation of JNK can be promoted. Its transcription factor c-Jun is formed, and c-Jun can bind to the transcriptional activator protein-1 (AP-1) site of many genes in the posterior region. Phosphorylation of p38 MAPK can also activate AP-1.

## Conclusions

In this study, we employed network pharmacology to explore and discuss multiple bezoar-mediated ischemic stroke pathways and treated targets. Our data suggest that bezoar blocks ischemic stroke through multiple signalling pathways. However, future research should provide experimental evidence, expand on the systemic level of bezoar in ischemic brain, and develop more effective drug delivery systems.

## Declarations

Ethics approval and consent to participate

Not applicable.

Consent for publication

Not applicable.

## Availability of data and material

The data used to support the findings of this study are available from the corresponding author upon request.

## Authors’ contributions

XD conceived and designed the project. CXL, BDR, ND, and CAC implemented the methods and conducted the analysis. XQW and FFC drafted the manuscript. QGW revised the manuscript. All authors read and approved the final manuscript.

## Acknowledgements

We greatly appreciate the support of the National Natural Science Foundation (grant numbers) and the Classical Prescription Basic Research Team of the Beijing University of Chinese Medicine.

## Abbreviations

PPI: Protein-protein interaction
TUDCA: Tauroursodeoxycholic acid
MCAO: Middle cerebral artery occlusion
TCMSP: Traditional Chinese Medicine Systems Pharmacology
OB: Oral bioavailability
PreOB: Predict oral bioavailability
PreDL: Predict drug-likeness
TTD: Therapeutic Target Database
OMIM: Online Mendelian Inheritance in Man
CTD: Comparative Toxicogenomics Database
DAVID: Database for Annotation, Visualization and Integrated Discovery
MAPK: Mitogen-activated protein kinase
ERK1/2: Extracellular signal-regulated kinase 1/2
JNK: c-Jun N-terminal kinase
PHD: Prolyl 4-hydroxylase domain
VEGF: Vascular endothelial growth factor
STAT3: Signal Transduction and Transcription Activator-3
ERK1/2: Extracellular signal-regulated kinase
NF-κB: Nuclear factor kappa B
AP-1: Activator protein-1

